# Development of DNA methylation-based epigenetic age predictors in loblolly pine (*Pinus taeda*)

**DOI:** 10.1101/2022.01.27.477887

**Authors:** Steven T. Gardner, Emily M. Bertucci, Randall Sutton, Andy Horcher, Doug Aubrey, Benjamin B. Parrott

## Abstract

Biological aging is connected to life history variation across ecological scales, as well as informing a basic understanding of age-related declines to organismal function. Altered DNA methylation dynamics are a conserved aspect of biological aging and have recently been modeled to predict chronological age among vertebrate species. In addition to their utility in estimating individual age, differences between chronological and predicted ages arise due to acceleration or deceleration of epigenetic aging, and these discrepancies are linked to disease risk and multiple life history traits. Although evidence suggests that patterns of DNA methylation can describe aging in plants, predictions with epigenetic clocks have yet to be performed. Here, we resolve the DNA methylome across CpG, CHG, and CHH-methylation contexts in the loblolly pine tree (*Pinus taeda*) and construct epigenetic clocks capable of predicting ages in this species within 8% of its lifespan. Although patterns of CHH methylation showed little association with age, both CpG and CHG methylation contexts were strongly associated with aging, largely becoming hypomethylated with age. Among age-associated loci were those in close proximity to malate dehydrogenase, NADH dehydrogenase, and 18S and 26S ribosomal RNA genes. This study reports one of the first epigenetic clocks in plants and demonstrates the universality of age-associated DNA methylation dynamics which can inform conservation and management practices, as well as our ecological and evolutionary understanding of biological aging in plants.

## Introduction

Alterations to the epigenome are a conserved hallmark of biological aging, and recent findings have demonstrated that age-associated DNA methylation patterns can be modeled to generate epigenetic age predictors capable of estimating chronological and biological age with unprecedented accuracy (Berdyshev, Korotaev, Boiarskikh, & Vaniushin, 1967; Christensen et al., 2009; Hannum et al., 2013; Richardson, 2003). In one of the first DNA methylation-based age predictors or “epigenetic clocks” developed by Horvath (2013), the methylation status of 353 cytosines predicts human chronological age with an error of ± 3.6 years. Epigenetic clocks have subsequently been developed in a variety of other mammalian (Horvath, 2013; Weidner et al., 2014), avian (Raddatz et al., 2021), and fish species (Anastasiadi & Piferrer, 2020; Bertucci, Mason, Rhodes, & Parrott, 2021; Mayne et al., 2020), and are currently being applied to biomedical and conservation problems, as well to questions regarding their relationship to the underlying biology of aging and senescence (Bertucci & Parrott, 2020; Kabacik, Horvath, Cohen, & Raj, 2018). However, whereas the phenomenon of epigenetic aging appears to be a conserved aspect of biological aging in vertebrates, age-associated changes to DNA methylation and their ability to predict chronological age in plants is relatively unexplored (Parrott & Bertucci, 2019).

DNA methylation refers to the covalent addition of a methyl group to the 5’ carbon of cytosine nucleotides (Jung & Pfeifer, 2015; Ng & Adrian, 1999; Suzuki & Bird, 2008), and although the functional consequences of these modifications vary across genomic context and taxonomic groups, DNA methylation is broadly associated with repressed transcriptional activity of genes and transposable elements through direct silencing and promotion of repressive chromatin states (Ng & Adrian, 1999; Zilberman, 2008). Similar to vertebrates, changes in DNA methylation levels in plants are observed with exposure to stress, age, and development (Dubrovina & Kiselev, 2016; Jiang et al., 2014; Law & Jacobsen, 2010; Probst & Mittelsten Scheid, 2015). However, compared to the distribution of cytosine methylation within vertebrate genomes (which almost exclusively occurs in CpG dinucleotides) DNA methylation in plant genomes is frequently observed within CpG, CHG and CHH contexts (where H = A, T, or C). Methylated cytosines located within gene bodies in plants primarily occurs in CpG contexts (Takuno, Ran, & Gaut, 2016), whereas CpG, CHG, and CHH methylation is typically found within highly repetitive genomic regions, potentially functioning to silence transposable element activity (Ausin et al., 2016; Slotkin & Martienssen, 2007). How these different sequence contexts might relate to age-associated DNA methylation patterning is not resolved.

Despite the ability of current epigenetic clocks to predict chronological age, discrepancies between epigenetic age and chronological age are observed and reflect variation in biological age (Horvath & Raj, 2018; Parrott & Bertucci, 2019; Xiao, Wang, & Kong, 2019). Accelerated epigenetic aging is associated with age related losses in organismal function and in humans, and predicts risk for age-associated disease and mortality (Levine et al., 2018; Perna et al., 2016; Zheng et al., 2016). In addition, epigenetic-to-chronological age discordance mirrors variation in life history traits (Anderson et al., 2021; Hamlat, Prather, Horvath, Belsky, & Epel, 2021). For example, birth weight, age and size at maturity, as well as the timing of reproductive senescence are all correlated to epigenetic age in humans, demonstrating intriguing links between aging processes and life history variation (Binder et al., 2018; Ryan et al., 2018; Simpkin et al., 2015). However, the links between the rate of epigenetic aging and variation in organismal function and life history traits (including those with potential commercial implications) are largely unexplored in plants.

Loblolly pine is a large tree with a broad geographic range spanning from southern New Jersey to eastern Texas, including parts of northern Florida (Baker & Langdon, 1990) and is capable of living for up to 275 years (Baker & Langdon, 1990). The loblolly pine is often commercially harvested for timber and is also used to diversify forest habitats, control erosion, and improve water quality (Baker & Langdon, 1990). Research and management practices aimed at improving growth and yield from loblolly pine stands often manipulate resource availability (Albaugh, Lee Allen, Dougherty, & Johnsen, 2004; Coyle, Aubrey, & Coleman, 2016; Fox, Lee Allen, Albaugh, Rubilar, & Carlson, 2007). However, the underlying biological mechanisms promoting optimal growth rates across variable environmental conditions remain unclear (Albaugh et al., 2004), and resolving age-associated methylation patterns in plants may aid in developing strategies for increasing traits such as leaf area and growth efficiency, which are linked to stand productivity (Dubrovina & Kiselev, 2016; Fox et al., 2007; Medlyn, Barrett, Landsberg, Sands, & Clement, 2003; Samuelson, Stokes, Cooksey, & McLemore III, 2001). Furthermore, a recent study examining DNA methylation in *Populus trichocarpa* reported correlations between epimutation and age (Shahryary et al., 2020; Yao, Schmitz, & Johannes, 2021), raising the possibility that these patterns could be used to predict other physiological traits such as plant growth (Hu, Morota, Rosa, & Gianola, 2015).

Here, we investigate age-associated DNA methylation patterns in *P. taeda* and evaluate their genomic distributions among differing methylation contexts. We then test differing modeling strategies to develop a novel epigenetic clock for *P. taeda*, which can be used to investigate the factors most important to growth, development, and tree aging. This study demonstrates the utility of epigenetic clocks in non-vertebrate models and indicates that age-associated DNA methylation may be a universal aspect of organismal aging.

## Materials and Methods

### Sample Collection

Between December 31, 2019 and January 14, 2020, cambium samples from standard coring procedures were obtained from 24 *P. taeda* individuals ranging from 1 to 119 years of age. This study was conducted at the United States Department of Energy’s Savannah River Site (Aiken, SC, USA), a National Environmental Research Park. The United States Department of Agriculture (USDA) Forest Service manages the natural resources of the Savannah River Site (Kilgo and Blake 2005). Using records from the Forest Service, we identified stands which were planted between 1 and 55 years prior and sampled three trees from each stand (1, 10, 19, 28, 37, 46, and 55 years old). Three additional trees of advanced age (82, 97, and 119 years old) were identified using core samples as planting records for older stands were unavailable. Cambium samples from adult trees (>1-year) were taken at breast height (1.37 meters) using a metal hole punch. Saplings (1-year old) were sampled at the base of the primary stem with a metal blade. All samples were immediately stored in RNAlater at -20°C until DNA extraction. Diameter at breast height was measured using a diameter tape and height was measured using a TruPulse 200x Rangefinder (Laser Technology, Inc., Centennial, CO).

### DNA extraction

DNA was extracted from cambium tissue using Qiagen’s DNeasy Plant Pro Kit (catalog # 69204, Qiagen, Hilden, Germany) following the manufacturer’s protocol with the addition of 100 μl of the Solution PS per sample due to high amounts of phenolic compounds in pine species. Briefly, samples were cut into small pieces using a sterile blade and then homogenized using a Mini-Beadbeater (BioSpec, Bartlesville, OK) for 4-8 min at 2000 oscillations/min. DNA was eluted in 50 μl of the supplied elution buffer, and the concentration and purity of DNA samples were assessed using a Qubit fluorometer 2.0 (Invitrogen, Carlsbad, CA) and Nanodrop spectrometer (Thermo-Scientific, Waltham, MA), respectively.

### Reduced Representation Bisulfite Sequencing Library Preparation

Reduced Representation Bisulfite Sequencing (RRBS) libraries were prepared using Diagenode’s Premium RRBS Kit (catalog #C02030032, Diagenode, Denville, NJ). Due to the occurance of CHG methylation in plant genomes, we adapted the protocol by digesting genomic DNA with the BsaWI restriction enzyme (Catalog #R0567S, New England BioLabs, Ipswich, MA) instead of MspI, as BsaWI cuts the recognition site W^CCGGW. Digestion efficiency of BsaWI in our samples was confirmed by visualizing digested genomic DNA from test samples using standard gel electrophoresis. For library preparations, 200 ng of genomic DNA from each sample was digested with 5 units/ng of BsaWI for 12 hr at 60°C, after which samples underwent a 20 min heat inactivation of the restriction enzyme at 80°C. To obtain sufficient library concentrations, two RRBS libraries were prepared for each sample. In the second library preparation, the protocol was further altered to add additional extension time in the final amplification (72°C for 1 min during cycling and 10 min during final extension.) Libraries were eluted in 22 μl of the supplied elution buffer and stored at -80°C until sequencing. Other than these alterations, the manufacturer’s protocol was followed exactly.

### RRBS Sequencing, Quality Control, and Alignment

RRBS libraries were assessed for concentration and fragment size distribution on a Fragment Analyzer (Advanced Analytical Technologies, Inc., Ames, IA) at the Georgia Genomics and Bioinformatics Core at the University of Georgia. Libraries were then pooled and sequenced single-end for 100 cycles on the Illumina NextSeq 2000 with 20% PhiX control added. Two library preparations for each sample were sequenced across five high-output flow cells and approximately 400 million reads from each flow cell were generated. The quality of the resulting reads was assessed using FastQC (v0.11.5) (Andrews, 2017). Reads were trimmed using TrimGalore! (v 0.4.5) to remove adapter sequences and low-quality reads (Phred score <25), using the –rrbs option for RRBS data. Trimmed reads were concatenated into a single file and aligned to a bisulfite converted index of the loblolly genome (Ptaeda2.0) using Bismark (v0.20.0) (Krueger & Andrews, 2011) allowing for one mismatch (option n -1). The subsequent alignments were sorted and indexed using SAMtools (v 0.1.19) (Li et al., 2009).

### File processing

The resulting Bam files were analyzed using the methylKit package (Akalin et al., 2012) in R (version 4.0.5) (2021). An average of 18,177,435 (+/-1,342,074) reads comprised each Bam file, and cytosines from each individual were divided into CpG, CHG, or CHH contexts using the read.context parameter in the processBismarkAln function of methylKit. To broadly assess age-associated DNA-methylation, while also developing an epigenetic clock, we filtered our data using two approaches. In the first approach, we filtered out cytosines covered by < 5x reads and not represented in < 80% of samples. To ensure comparability across CpG, CHG, and CHH contexts, cytosines on opposite strands were not merged (destrand = FALSE). This data set is considered our “exploratory” data set, in which general age-associated patterns could be resolved. In the second approach, we increased coverage requirements (≥ 10x) to yield a dataset that could be used to construct epigenetic age predictors.

### Exploratory data set

The resulting exploratory data set was analyzed using the files generated for CpG, CHG, and CHH cytosine-methylation contexts. We filtered out cytosines displaying zero or near-zero variance in methylation status across samples using the nearZeroVar function from the Classification And Regression Training (caret) package (Kuhn, 2015) in R. Following filtering of invariant cytosines, we performed Spearman correlations between the methylation status of each cytosine and chronological age using the corr.test function from the psych package (Revelle, 2019) in R. Given the exploratory nature of the downstream analyses, we considered cytosines with an absolute correlation coefficient greater than 0.5 (|R| > 0.5) as being “age-associated”, and those with p-values less than 0.05 (*P* < 0.05) following an FDR correction for multiple comparisons as “significantly correlated” with age.

To assess genomic characteristics of age-associated and significantly correlated CpG, CHG, and CHH cytosines, we classified the genomic locations of all covered cytosines according to genomic context within the *P. taeda* genome using the GenomicRanges package (Lawrence et al., 2013). We first used CpG plot (Madeira et al., 2019) to identify CpG islands (CGIs; parameters: widow = 100, min len = 200, minoe = 0.6, minpc = 50). We then used CGI coordinates to generate coordinates for shore regions (± 2000 bp from CGIs), and shelf regions (± 2000-4000 bp from CGIs) using Bedtools and Samtools (H. Li et al., 2009; Quinlan & Hall, 2010). All remaining sites not falling in island, shore, or shelf regions were characterized as open sea regions. Enrichment above background within each methylation context was performed with binomial tests using all cytosines following invariant filtering within each context (21,566, 25,501, and 33,151 CpG, CHG, and CHH cytosines, respectively) as the background levels. To identify genes in close proximity to cytosines according to age-associated methylation patterns, we performed BLAST searches on 400 bp regions (200bp upstream and 200 bp downstream) centered around each of the 35 cytosines from CpG and CHG contexts with the greatest correlations with age.

### Clock data set

Our exploratory analyses indicated that the methylation status of CHH cytosines was not associated with age; thus, we only generated epigenetic clocks using CpG and CHG cytosines. We first removed any invariant sites using the nearZeroVar function from the caret package (Kuhn, 2015) and performed imputation on missing data using a K-nearest neighbor (KNN) approach in the impute package (Hastie et al. 2021) in R. We set k=2 as each age group among our samples was only represented by a maximum of three samples. We then split our data into a training data set (n = 19; distributed among ages 1, 10, 19, 28, 37, 46, 55, and 97 years old), and a test data set (n = 5; ages: 1, 28, 55, 82, and 119 years old). Individuals comprising the test set were selected by dividing the age range (1 – 119 years) into 5, then choosing 5 individuals with ages separated by approximately 24 years.

### Elastic net clocks

We trained elastic net models for CpG and CHG methylated cytosines using the glmnet package (Friedman, Hastie, & Tibshirani, 2010) in R. We used an elastic net penalized regression model (alpha = 0.5, family = gaussian) to select CpG-and CHG-methylated cytosines and assign penalties to individual model coefficients using the subset of CpG-loci and CHG-loci. We used a leave-one-out cross validation approach (nfold = 19) to select the optimal lambda value (value resulting in minimum mean error) for the training set model (n = 19 samples). We then used our training set model to predict ages of the individuals in our test set (n = 5). Following the generation of our elastic net epigenetic clock, we performed BLAST searches of 400 bp regions centered around each clock cytosine and used GenomicRanges to determine the genomic contexts for these sites (genes and CpG islands, shores, shelves, or open seas).

### Pearson clocks

To construct linear models capable of predicting age, we identified the top 5 and 10 cytosines with the greatest Pearson correlation with age (|R| > 0.5) from our training set using the corr.test function (using FDR correction, alpha = 0.05) from the psych package. We generated our linear models using the lm function in R, and then used them to predict the ages of the individuals in our test data set. Following the generation of our linear model epigenetic clocks, we performed BLAST searches of 400 bp regions centered around clock cytosines and used GenomicRanges to determine genomic context (genes and CpG islands, shores, shelves, or open seas).

To compare performance of our elastic net and Pearson clocks, we used multiple regression to compare the mean absolute errors (MAE) generated when using each clock to predict ages of trees from our test set. We used clock type as a factor and chronological age as a covariate and corrected resulting *P* values using Bonferroni correction for multiple comparisons. Additionally, overfit of each clock to our training set was evaluated by comparing the MAE for training and test sets from each clock using t-tests.

## Results

### Characterizing age-related cytosine methylation

Among the CpGs analyzed, 533 (2.47%) displayed age-associated methylation, with 207 CpGs having positive correlations (max R = 0.84) and 326 having negative correlations (min R = -0.79) (Figure 1). Following FDR correction for multiple comparisons, 10 CpGs (9 negative and 1 positive) showed significant correlations with age. When examining locations of our age-associated cytosines with respect to genomic contexts, CpG-methylated cytosines were enriched in island (*P* = 4.85e-10) and shore regions (*P* = 0.038), while being depleted in shelf (*P* = 9e-04) and open-sea regions (*P* = 2.20e-06) (Figure 2). Among the age-associated CpG-methylated sites found in islands, many became hypomethylated with age (Table 1). Whereas this pattern was also observed for CpGs in shore and open-seas, CpGs in shelf regions were more evenly split between hypo-and hypermethylation with increasing age. Malate dehydrogenase (mdh), as well as 18S and 26S ribosomal genes of related pine species, were among the genes that a few of our methylated CpG-containing 400bp regions mapped to (Supplemental Table 1).

**Figure 1.**
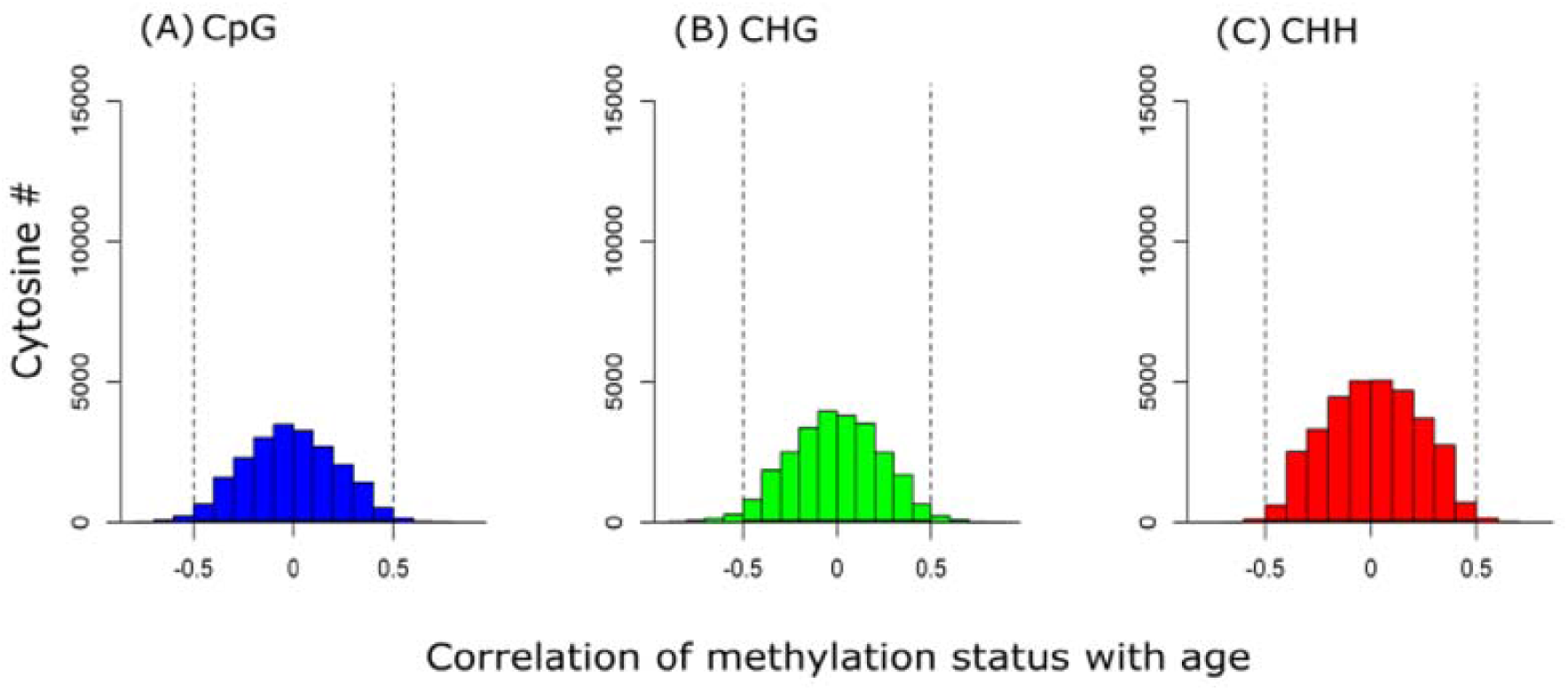
Distributions of Spearman correlation coefficients of cytosine methylation status with age following filtering of invariant sites across CpG, CHG, and CHH methylation contexts from 24 Loblolly pine trees of differing ages. (A) Of 21,567 CpGs analyzed, 533 (2.5%) showed correlations between methylation status and age of R > |0.5|. (B) Of 25,501 CHGs analyzed, 869 (3.4%) displayed correlations between methylation status and age of R > |0.5|. (C) Of 33,151 CHHs analyzed, 308 (0.93%) showed correlations between age and methylation status of R > |0.5|.

**Figure 2.**
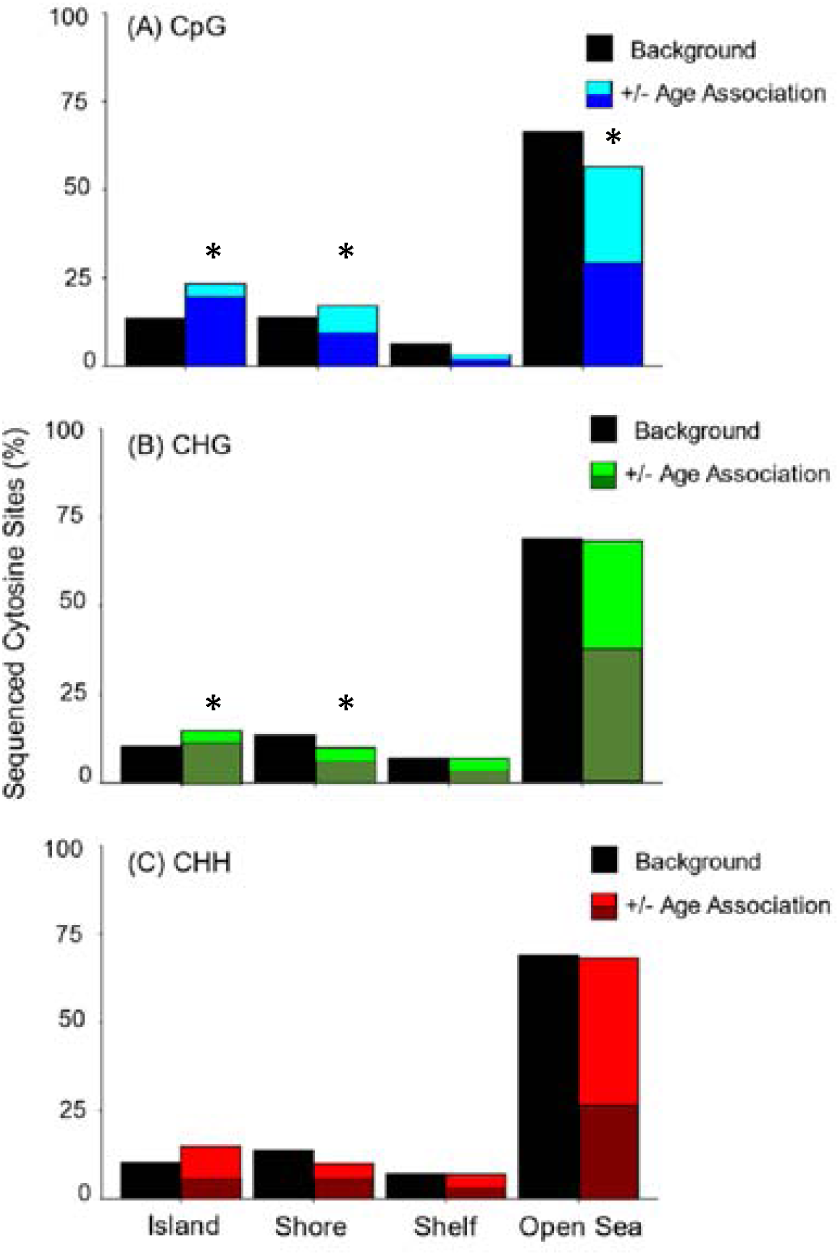
Enrichment and depletion of age-related DNA methylation within genomic regions with varying CpG densities. Cytosines for which methylation status was associated with age (|R > 0.5|) were compared against all RBSS-captured cytosines (background). (A) Age-associated CpG methylation was enriched in CpG island and shore regions and were depleted in CpG shelf and open-sea regions. (B) Age-associated CHG methylation was enriched in CpG islands and depleted in shore regions. (C) There were no differences in the distributions of age-associated CHH-methylated cytosines compared to background CHH methylated cytosine levels (Asterisks indicate *P* < 0.05).

**Table 1.** Distributions of hypo- (-) and hypermethylated (+) cytosines with age among genomic regions

| Context | Cytosine subset | Island Cytosines |  | Shore Cytosines |  | Shelf Cytosines |  | Open sea cytosines |  |
| --- | --- | --- | --- | --- | --- | --- | --- | --- | --- |
|  |  | - | + | - | + | - | + | - | + |
| CpG | Age-associated | 111 | 14 | 56 | 35 | 7 | 9 | 152 | 149 |
|  | Significantly correlated | 2 | 0 | 2 | 1 | 0 | 0 | 4 | 1 |
| CHG | Age-associated | 105 | 24 | 57 | 30 | 33 | 27 | 325 | 269 |
|  | Significantly correlated | 21 | 0 | 5 | 1 | 1 | 0 | 33 | 9 |
| CHH | Age-associated | 7 | 14 | 20 | 14 | 11 | 12 | 84 | 147 |

For CHG methylation, 869 cytosines (3.41%) were age associated, with 350 displaying positive correlations (max R = 0.87) and 519 negatively correlated with age (min R = -0.89; Figure 1). Following FDR corrections, 70 of these 869 age-associated sites (60 hypomethylated and 10 hypermethylated) showed significant (*P* < 0.05) correlations with age. Compared to background levels, age-associated CHG cytosines were enriched in island regions (*P* = 3.62e-05) and were depleted in shores (*P* = 1.3e-03) (Figure 2). The majority of CHG-methylated cytosines were hypomethylated with age, regardless of genomic context (Table 1). One of the top 35 CHG age-associated cytosine sites mapped to an apparent homolog of mdh and became hypermethylated with age.

Within CHH-methylation contexts, 308 (0.093%) cytosines were found to be age-associated, with 187 displaying a positive correlation with age (max R = 76) and 121 negatively correlated to age (min R = -0.71; Figure 1). However, following FDR corrections, no cytosines were significantly (*P* < 0.05) correlated with aging, and compared to background levels, there were no differences in the distributions of age-associated CHH-methylated cytosines based on genomic context (*P* > 0.05). CHH-methylated cytosines found in island regions showed a trend of being mostly hypermethylated (67%) with age (Table 1), while CHHs found in shore regions were mostly hypomethylated (59%) with age. Shelf-region CHHs were more evenly split between hypo and hypermethylation with age (48% vs 52%), while most of the open-sea region CHH cytosines showed hypermethylation with age (63%) (Figure 2).

### Construction of epigenetic clocks capable of predicting age

Individual age predictors were initially constructed using elastic net regularized regression approaches using either CpG methylation, CHG methylation, or a combination of CpG and CHG methylation. All three elastic net clocks predicted ages of trees from our training set within 0.62 years, and predicted ages were highly correlated with chronological ages for these trees (R^2^ = 0.99) (Table 2). However, the ability of these models to predict chronological ages of trees in the test set were slightly lower and R^2^ values ranged from 0.79 to 0.90, with the CHG model outperforming the CpG model, and the CpG and CHG combined model (Table 3) performing the best.

**Table 2.** Performance of elastic net models when predicting ages of trees from training and test

| Elastic Net |  |  |  |  |  |  |  |  |
| --- | --- | --- | --- | --- | --- | --- | --- | --- |
| Context | # Initial sites | # Filtered sites | # Clock sites | Train R2 | Train MAE (years) | Test R2 | Test MAE (years) | Overfit? (t-test p-value) |
| CpG | 26520 | 18844 | 35 | 0.9998 | 0.62 (+/- 0.08) | 0.79 | 22.68 (+/- 12.02) | No (0.14) |
| CHG | 30728 | 22263 | 50 | 0.9999 | 0.56 (+/- 0.10) | 0.843 | 25.00 (+/- 11.52) | No (0.10) |
| CpG+CHG | 57248 | 41107 | 47 | 0.9999 | 0.53 (+/- 0.10) | 0.899 | 22.53 (+/- 9.99) | No (0.09) |

**Table 3.**
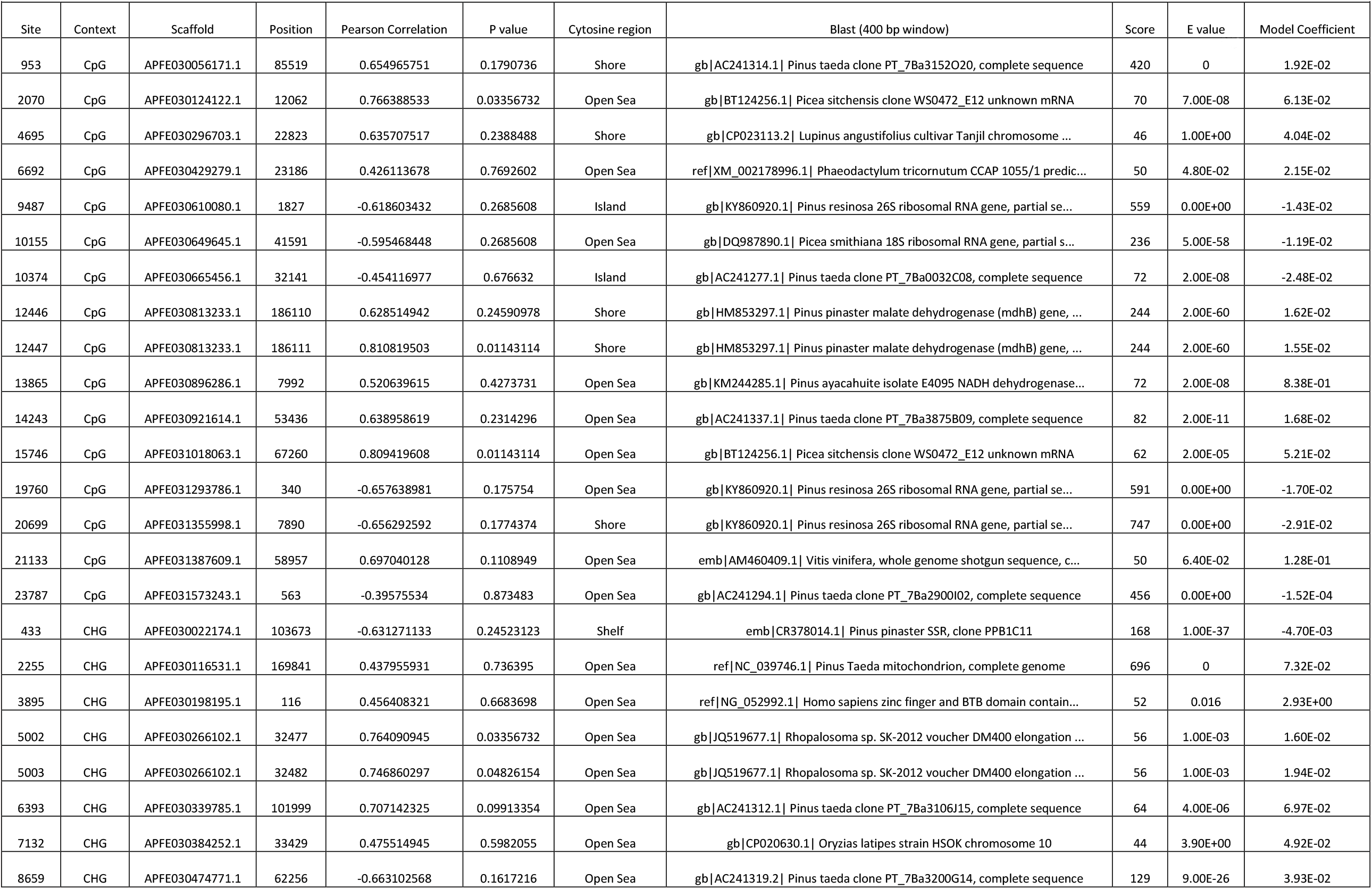
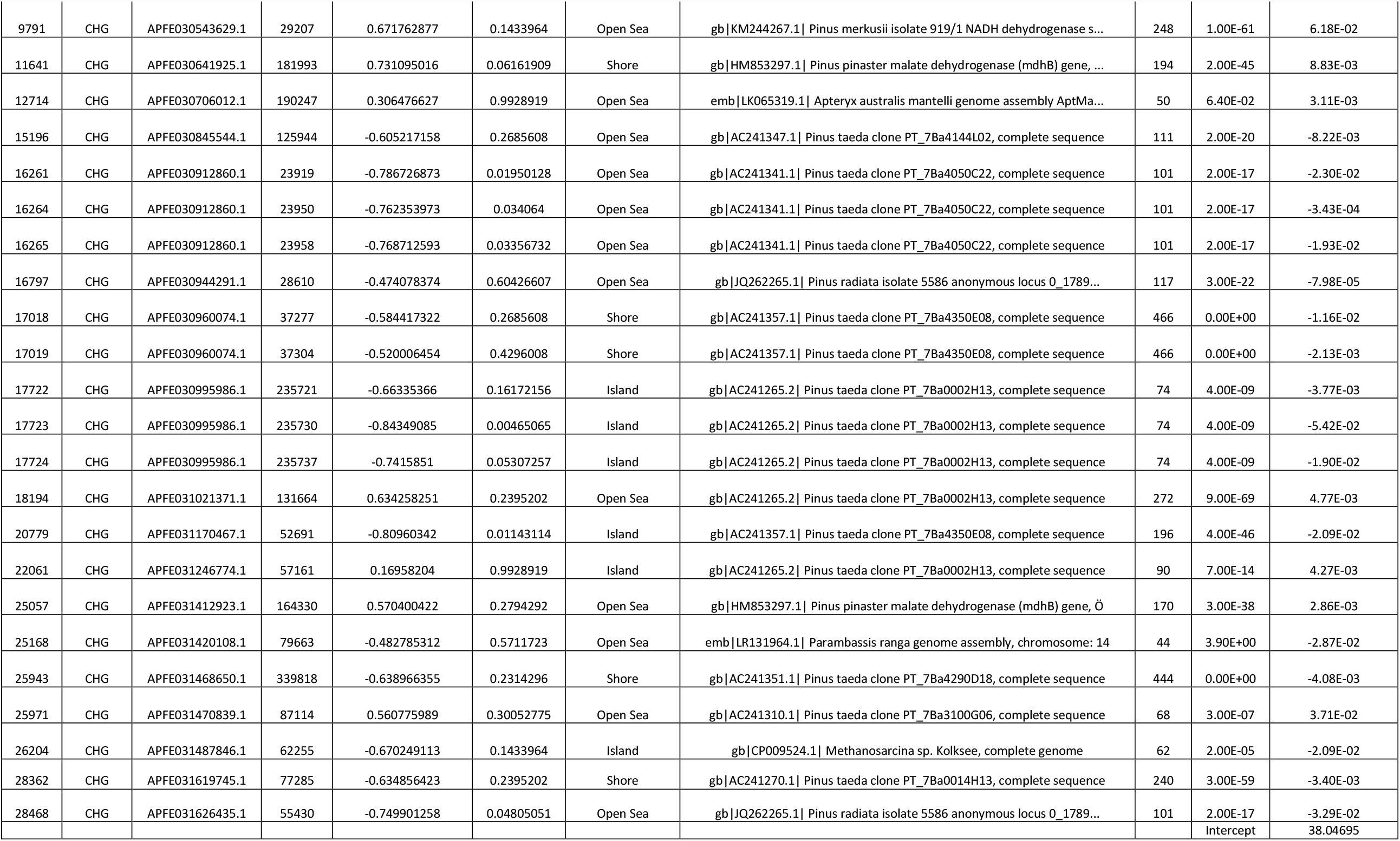
Combined CpG and CHG elastic net model

In addition to our elastic net models, we manually constructed epigenetic clocks for each methylation context (CpG, CHG, and CpG + CHG) using cytosines with the greatest Pearson correlation coefficients. When the top 5 Pearson age-correlated cytosines were selected, ages were predicted within 3.91 years and were highly correlated to chronological age (R^2^ = 0.89 to 0.95) (Table 4). Predictions from the combined model (Table 5) again showed the highest correlations with chronological age. Correlations (R^2^) between predicted and chronological ages ranged from 0.83 when using only CpG-methylated cytosines to 0.9 for our model using a combination of CpG and CHG cytosines. We then constructed models using the top 10 Pearson age-corelated cytosines. Relative to the models using the top 5 correlated cytosines, these models performed better on the training set (R^2^ = 0.94 to 0.98); however, performance suffered on the test set, R^2^ values ranging from 0.03 when using only CpG-methylated cytosines to 0.87 when using both CpG and CHG-methylated cytosines.

**Table 4.** Performance of Pearson models when predicting ages of trees from training and test sets

| Pearson Model (5 cytosines) |  |  |  |  |  |  |  |
| --- | --- | --- | --- | --- | --- | --- | --- |
| Context | # Initial sites | # Filtered sites | Train R2 | Train MAE (years) | Test R2 | Test MAE (years) | Overfit? (t-test p-value) |
| CpG | 26520 | 18844 | 0.898 | 5.48 (+/- 1.13) | 0.829 | 24.02 (+/- 9.9) | No (0.13) |
| CHG | 30728 | 22263 | 0.911 | 5.84 (+/- 0.81) | 0.873 | 20.86 (+/- 9.10) | No (0.17) |
| CpG+CHG | 57248 | 41107 | 0.952 | 3.91 (+/- 0.72) | 0.9 | 22.18 (+/- 9.41) | No (0.12) |
| Pearson Model (10 cytosines) |  |  |  |  |  |  |  |
| Context | # Initial sites | # Filtered sites | Train R2 | Train MAE (years) | Test R2 | Test MAE (years) |  |
| CpG | 26520 | 18844 | 0.943 | 4.07 (+/-0.84) | 0.03 | 38.14 (+/- 19.03) | No (0.15) |
| CHG | 30728 | 22263 | 0.972 | 3.28 (+/- 0.45) | 0.58 | 22.08 (+/- 13.41) | No (0.23) |
| CpG+CHG | 57248 | 41107 | 0.979 | 2.59 (+/- 0.5) | 0.87 | 21.12 (+/- 9.01) | No (0.11) |

**Table 5.** Combined CpG and Pearson models (5 and 10 cytosines)

| Site | Context | Scaffold | Position | Pearson Correlation | P value | Cytosine region | Blast (400 bp window) | Score | E value | Top 5 Model Coefficient | Top 10 Model Coefficient | Intercept |
| --- | --- | --- | --- | --- | --- | --- | --- | --- | --- | --- | --- | --- |
| 21133 | CpG | APFE031387609.1 | 58957 | 0.697040128 | 0.111 | Open Sea | emb\|AM460409.1\| Vitis vinifera, whole genome shotgun sequence | 50 | 6.40E-02 | 0.242 | 0.32154 |  |
| 17723 | CHG | APFE030995986.1 | 235730 | -0.84349085 | 0.004 | Island | gb\|AC241265.2\| Pinus taeda clone PT_7Ba0002H13, complete sequence | 88 | 3.00E-13 | -0.231 | -0.21276 |  |
| 2070 | CpG | APFE030124122.1 | 12062 | 0.766388533 | 0.033 | Open Sea | gb\|BT124256.1\| Picea sitchensis clone WS0472_E12 unknown mRNA | 70 | 7.00E-08 | 0.223 | 0.20857 |  |
| 15746 | CPG | APFE031018063.1 | 67260 | 0.809419608 | 0.011 | Open Sea | gb\|BT124256.1\| Picea sitchensis clone WS0472_E12 unknown mRNA | 62 | 2.00E-05 | 0.009 | -0.15923 |  |
| 6393 | CHG | APFE030339785.1 | 101999 | 0.707142325 | 0.099 | Open Sea | gb\|AC241312.1\| Pinus taeda clone PT_7Ba3106J15, complete sequence | 64 | 4.00E-06 | 0.281 | 0.37791 | Top 5 : 30.67 |
| 12447 | CpG | APFE030813233.1_ | 186111_ | 0.810819503 | 0.011 | Shore | PREDICTED: Oryzias melastigma trace amine-associated receptor_ | 44_ | 3.9_ |  | 0.06751 |  |
| 7132 | CHG | APFE030384252.1 | 33429 | 0.807122 | 0.113 | Open Sea | gb\|CP020630.1\| Oryzias latipes strain HSOK chromosome 10 | 44 | 3.90E+00 |  | -0.44427 |  |
| 16261 | CHG | APFE030912860.1_ | 23919_ | -<br>0.786726873 | 0.020 | Open Sea | gb\|AC241341.1\| Pinus taeda clone PT_7Ba4050C22, complete sequence_ | 101_ | 2.00E-17_ |  | -0.07754 |  |
| 20779 | CHG | APFE031170467.1_ | 52691_ | -0.80960342 | 0.011 | Island | gb\|AC241357.1\| Pinus taeda clone PT_7Ba4350E08, complete sequence_ | 196_ | 4.00E-46_ |  | 0.05243 |  |
| 5002 | CHG | APFE030266102.1 | 32477 | 0.805784 | 0.113 | Open Sea | gb\|JQ519677.1\| Rhopalosoma sp. SK-2012 voucher DM400 elongation ... | 56 | 1.00E-03 |  | 0.05348 | Top 10: 29.79 |

When comparing predictive performance among our elastic net and Pearson clocks, MAE of predicted ages for trees from our test set ranged from 20.86 to 38.14 years (7.58-13.9% of the total lifespan for *P. taeda*). Although MAE for our test set did not differ based on clock type (*P* > 0.05), MAE of predicted ages increased by 0.43 (± 0.06) years with each chronological year increase in age for trees in our test set (*t*_35_ = 7.04, *P* = 1.02e-06) (Figures 3 and 4). To assess the potential overfit of our models, we compared the MAE of the training sets to those of test sets. None of our models were overfit to our training set (*P* > 0.05) (Tables 2 and 4).

**Figure 3.**
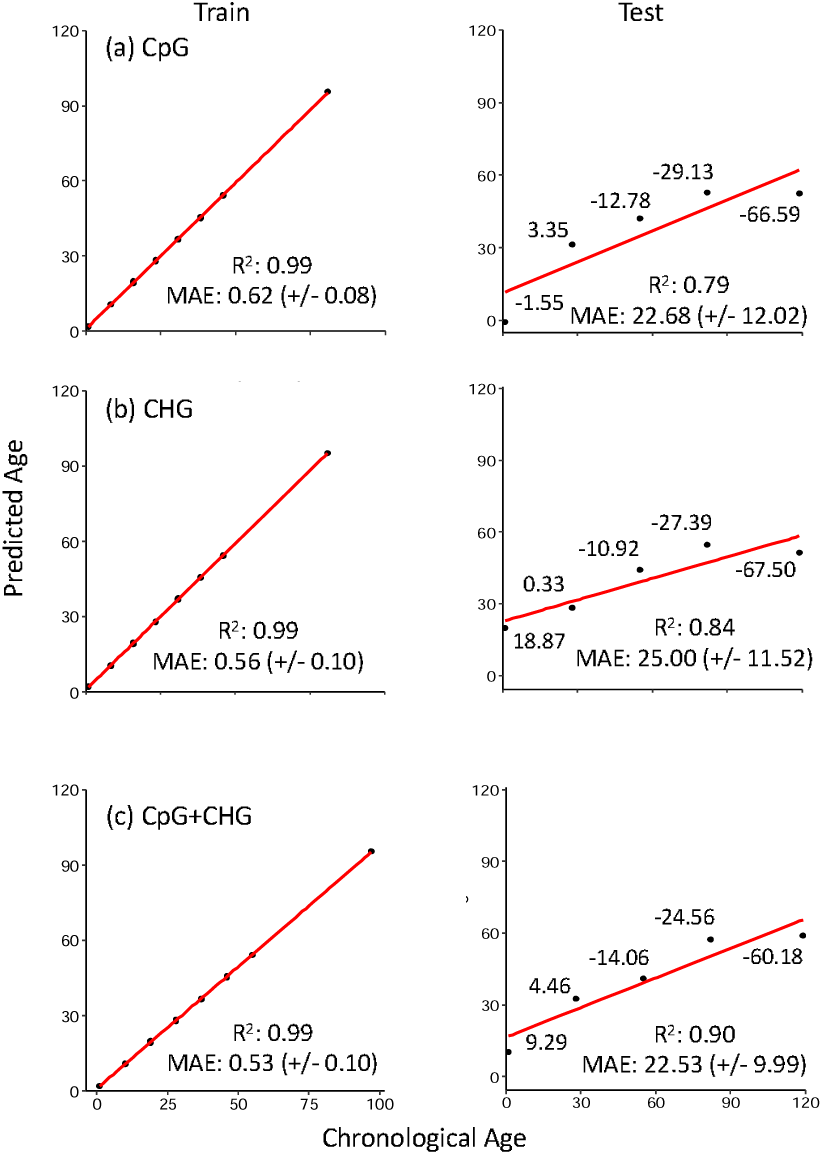
Elastic net models used to predict chronological age in *P. taeda* for both training and test subsets. Clock accuracy is measured by the Pearson’s correlation coefficient and precision by the mean absolute error (MAE) (+/- standard error). (A) 35 CpGs out of 18,844 CpGs following invariant filtering were selected in our elastic net model. (B) 50 CHGs out of 22,263 CHGs were selected and incorporated into the clock. (C) Combining CpG and CHG datasets, there were 41,107 cytosines following invariant filtering, with 47 incorporated into the clock. Values of individual datapoints represent the error between predicted ages from our models compared to chronological ages among test individuals, with positive values indicating overestimation and negative values indicating underestimation of chronological age.

**Figure 4.**
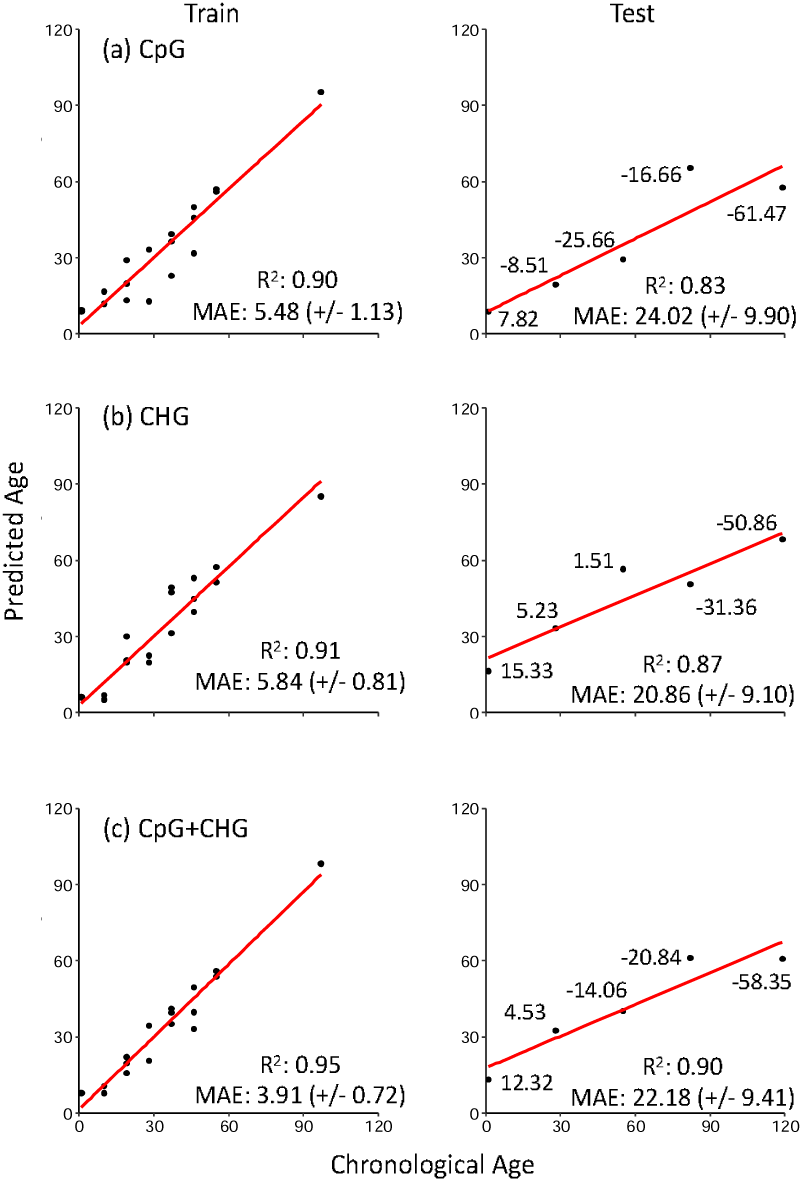
Pearson models incorporating the five cytosines with the strongest age-associated methylation patterns predict chronological ages in *P. taeda* for both training and test subsets for CpG, CHG, and combined (CpG+CHG) models. Clock accuracy is measured by the Pearson’s correlation coefficient and precision by the mean absolute error (MAE) (+/- standard error). Values of individual datapoints represent the error between predicted ages from our models compared to chronological ages among our test individuals, with positive values indicating overestimation and negative values indicating underestimation of chronological age.

We next used BLAST to assess if clock cytosines were proximal to specific genes. Among the loci that mapped to known genes were mdh, NADH dehydrogenase, and rRNA (18S and 26S). These loci were found in both methylation contexts (CpG and CHG) among our clocks, and were distributed among all genomic contexts (islands, shores, shelves, and open seas). Both mdh and NADH dehydrogenase sites became hypermethylated with age, and and sites mapping to rRNA became hypomethylated with age (similar to patterns we observed in our exploratory data set). Within our combined CpG+CHG elastic net clock 10 of 47 cytosines returned hits based on homology to these genes (Table 5). Among the 4 cytosines that mapped to mdh, the 2 CpG loci were located next to each other on the same scaffold, and both were in shore regions.

The remaining 2 CHG loci that mapped to mdh were in shore and open sea regions on different scaffolds. Similar to mdh, the 2 cytosines that mapped to NADH dehydrogenase were also located on different scaffolds, 1 being a CpG and located in an open sea region and the other a CHG found in an open sea region. Cytosines in rRNA genes (18S and 26S) were CpG-methylated, with the 18S cytosine being found in an open sea region and the 26S cytosines (3) found in island, shore, and open sea regions on differing scaffolds of the *P*.*taeda* genome, respectively. For annotations of cytosines from CpG and CHG clocks, see Supplemental Tables 3-6.

## Discussion

We demonstrate the first use of methylation patterns to generate epigenetic clocks for *P. taeda*, which are capable of predicting chronological ages within ∼8% of the total lifespan (275 years) of this species. Although there were no differences in test set MAE among our clocks, predicted ages from clocks combining methylation in both contexts showed the highest correlations with age, indicating that methylation in both contexts is useful when predicting ages for *P. taeda* and potentially other plant species. Although age-associated CpG and CHG methylation was distributed across genomic regions, we observed an enrichment of age-associated CpG methylation within island and shore regions, and CHG methylation in island regions. Within both methylation contexts, age-associated methylation mostly resulted in hypomethylation, regardless of genomic (island, shore, shelf, or open sea) context. In regard to potential functional implications, only CpG-methylation has been previously associated with gene expression in *P. taeda* (Takuno et al., 2016). As increased CpG methylation at promoter sequences (commonly associated with CpG island regions) is associated with reduced expression of downstream genes (Gehring & Henikoff, 2007), hypomethylation at many island-region sites may indicate that expression of specific genes increases with age in *P. taeda*. In contrast to CpG-methylation, CHG-methylation is associated with gene splicing (Chaudhary, Jabre, & Syed, 2021; Zhang, Lang, & Zhu, 2018). Alterations in CHG-methylation status can affect the proper splicing of genes (Chaudhary et al., 2021; Zhang et al., 2018) which can have deleterious consequences (Ong-Abdullah et al., 2015). When comparing age-related changes in methylation status among cytosines from both CpG and CHG contexts mapping to known genes in our BLAST searches, gains and losses were consistent for sites mapping to similar genes for both contexts. Thus, our findings taken together with current functional understanding indicates a possible link between epigenetic aging and increases in gene expression and splicing variation with age. Compared to CpG and CHG-methylation, CHH-methylation in plants has largely been associated with transposable element silencing (Dubin et al., 2015), and rates of epimutations in transposable element regions are generally low (van der Graaf et al., 2015). Thus, the stability of CHH methylation through development could explain the poor associations between CHH methylation and aging observed in the current study.

Although the majority of cytosines in our analyses did not map to annotated genes, the expression of critical genes involved in conserved molecular pathways that regulate life-history traits has been observed to influence multiple physiological processes (Partridge & Gems, 2002; Perls et al., 2002; Flatt & Partridge, 2018). Among the genes proximal to several of our age-associated CpG and CHG-methylated sites in the exploratory and clock data sets were malate dehydrogenase (mdh), NADH dehydrogenase, and 18S and 26S rRNA. The expression of mdh, a multi-subunit metabolic enzyme in many organisms including plants (Longo & Scandalios, 1969; Yudina, 2012), has been linked to respiration and CO_2_ assimilation rates, and normal plant growth and development (Tomaz et al., 2010). Expression has been shown to increase with age, which is postulated to maintain aerobic metabolism (Sharma & Patnaik, 1982). NADH dehydrogenase, another important metabolic enzyme involved in the electron transport chain (Weiss, Friedrich, Hofhaus, & Preis, 1992), is also associated with plant growth and development (Sweetman et al., 2019). Proper subunit splicing of this enzyme is essential for its function (Bonen, 2008; Malek & Knoop, 1998), with defective splicing leading to decreased activity of the electron transport chain, slower growth, and delayed germination and phenotypic development (Hsieh et al., 2015). Expression and activity of rRNA have been shown to increase with increased metabolic demands (Russell & Zomerdijk, 2005), also affecting growth, cell adaptation, stress responses, and cell proliferation (Russell & Zomerdijk, 2005). Increased methylation near rRNA promoter regions greatly reduces expression (Ghoshal et al., 2004; Zatsepina et al., 1993); therefore, the loss of methylation we observed for cytosines proximal to 18S and 26S rRNA genes in island and shore regions coupled with the methylation gains we observed for CpG-methylated cytosines near mdh and NADH dehydrogenase in shore and open sea regions may indicate expression of these genes increases to match increasing metabolic demands as *P. taeda* trees age. If so, an increased requirement for proper splicing of NADH and mdh would be needed (Gendrel, Lippman, Yordan, Colot, & Martienssen, 2002; Jeddeloh, Stokes, & Richards, 1999), which might explain the increased CHG-methylation near these genes with age; although these predictions require further evaluation.

Beyond their ability to predict chronological age, the epigenetic clocks developed here are likely to find utility in forest management applications and addressing questions surrounding the basic biology of aging. The directionality and magnitude of epigenetic-to-chronological age discordance in humans and other vertebrates correlates to variation in life history traits (Anderson et al., 2020; Hamlat et al., 2021; Ryan, 2021). Variation in life history traits can have consequences for stand productivity (Kellner & Swihart, 2016; Schulze, 2003). If epigenetic-to-chronological age discordance, especially in early life, is connected to variation in tree growth, productivity, wood quality, and/or responses to disturbances, epigenetic clocks might inform breeding programs and could be used in evaluating efficiencies of experimental manipulations to increase early-life growth and subsequent wood yield from managed *P. taeda* stands (Baker & Langdon, 1990; Clason, 1989). Age-associated variation in DNA methylation patterns is also associated with species-specific lifespans in vertebrates (Mayne, Berry, Davies, Farley, & Jarman, 2019). The findings presented here raise the possibility that age-related DNA methylation patterns might be used to estimate lifespan and yield insights into the underlying genetic and epigenetic determinants across different plant species.

A limitation of this study was the relatively small number of individual trees analyzed. The *P. taeda* genome, like that of many tree species, is quite large (20 billion base pairs) (Neale et al., 2014). As a result, resolving patterns of genomic methylation are sequencing intensive, and more cost-efficient technical approaches (e.g. amplicon sequencing, bait capture techniques) are likely needed if measures of epigenetic age are to be scaled to stand and population applications (Meek & Larson, 2019). Due to the age distributions of our sampled trees, age predictions for our individuals older than 55 years of age were poor, which may have contributed to the greater MAE from our clocks compared to those predicting ages for many vertebrate species (Bertucci et al., 2021; Bors et al., 2021; Lemaître et al., 2020; Meer, Podolskiy, Tyshkovskiy, & Gladyshev, 2018; Stubbs et al., 2017). As we initially removed sites not present in at least 80% of our individuals and many of our trees were within stand ages (Cunningham, n.d.; Li et al., 1999; Stiff & Stansfield, 2003), additional sampling of older trees could allow for retention of higher numbers of cytosine sites whose methylation status remains dynamic in later life, allowing for more accurate age predictions of older trees in further clock studies. Despite these limitations, we clearly demonstrate the relationship between DNA methylation and chronological age in a long-lived tree species and indicate that alterations in DNA methylation may be a universal aspect of aging across the tree of life.

## Acknowledgements

We would like to thank Samantha Bock for her assistance with bioinformatic work, as well as the members of the Parrott lab at SREL for their feedback pertaining to the results of this study. This study was supported in part by the USDA Forest Service-Savannah River, under Interagency DE-EM0003622 with the U.S. Department of Energy, the National Science Foundation (Award #2026210, BBP), and the U.S. Department of Energy Office of Environmental Management under award number DE-EM0004391 to the University of Georgia Research Foundation.

## Disclaimer

This report was prepared as an account of work sponsored by an agency of the United States Government. Neither the United States Government nor any agency thereof, nor any of their employees, makes any warranty, express or implied, or assumes any legal liability or responsibility for the accuracy, completeness, or usefulness of any information, apparatus, product, or process disclosed, or represents that its use would not infringe privately owned rights. Reference herein to any specific commercial product, process, or service by tradename, trademark, manufacturer, or otherwise does not necessarily constitute or imply its endorsement, recommendation, or favoring by the United States Government or any agency thereof. The views and opinions of authors expressed herein do not necessarily state or reflect those of the United States Government or any agency thereof.

## Data accessibility

Data used to support the claims in this manuscript will be made publicly available in the NCBI Sequence Read Archive (SRA) upon acceptance for publication. Scripts used to analyze data will be publicly available at https://github.com/stg0015/Loblolly-scripts.

## Author contributions

E.M.B., B.B.P, D.P.A., and A.H. developed the concept for the paper. E.M.B. and R.S. collected the samples. E.M.B. prepared the libraries and assisted S.G. with data analysis. S.G. analyzed the data and wrote the paper while being assisted by E.M.B. and B.B.P.

## References

Akalin, A., Kormaksson, M., Li, S., Garrett-Bakelman, F. E., Figueroa, M. E., Melnick, A., & Mason, C. E. (2012). methylKit: A comprehensive R package for the analysis of genome-wide DNA methylation profiles. Genome Biology, 13(10), R87. doi: 10.1186/gb-2012-13-10-r87

Albaugh, T. J., Lee Allen, H., Dougherty, P. M., & Johnsen, K. H. (2004). Long term growth responses of loblolly pine to optimal nutrient and water resource availability. Forest Ecology and Management, 192(1), 3–19. doi: 10.1016/j.foreco.2004.01.002

Anastasiadi, D., & Piferrer, F. (2020). A clockwork fish: Age prediction using DNA methylation-based biomarkers in the European seabass. Molecular Ecology Resources, 20(2), 387–397. doi: 10.1111/1755-0998.13111

Anderson, J. A., Johnston, R. A., Lea, A. J., Campos, F. A., Voyles, T. N., Akinyi, M. Y., … Tung, J. (2020). The costs of competition: High social status males experience accelerated epigenetic aging in wild baboons (p. 2020.02.22.961052). doi: 10.1101/2020.02.22.961052

Anderson, J. A., Johnston, R. A., Lea, A. J., Campos, F. A., Voyles, T. N., Akinyi, M. Y., … Tung, J. (2021). High social status males experience accelerated epigenetic aging in wild baboons. ELife, 10, e66128. doi: 10.7554/eLife.66128

Andrews, S. (2017). FastQC: a quality control tool for high throughput sequence data. 2010.

Ausin, I., Feng, S., Yu, C., Liu, W., Kuo, H. Y., Jacobsen, E. L., … Wang, H. (2016). DNA methylome of the 20-gigabase Norway spruce genome. Proceedings of the National Academy of Sciences, 113(50), E8106–E8113. doi: 10.1073/pnas.1618019113

Baker, J. B., & Langdon, O. G. (1990). Pinus taeda L. Loblolly pine. Silvics of North America, 1, 497–512.

Berdyshev, G. D., Korotaev, G. K., Boiarskikh, G. V., & Vaniushin, B. F. (1967). Nucleotide composition of DNA and RNA from somatic tissues of humpback and its changes during spawning. Biokhimiia (Moscow, Russia), 32(5), 988–993.

Bertucci, E. M., Mason, M. W., Rhodes, O. E., & Parrott, B. B. (2021). The aging DNA methylome reveals environment-by-aging interactions in a model teleost (p. 2021.03.01.433371). doi: 10.1101/2021.03.01.433371

Bertucci, E. M., & Parrott, B. B. (2020). Is CpG Density the Link between Epigenetic Aging and Lifespan? Trends in Genetics, 36(10), 725–727. doi: 10.1016/j.tig.2020.06.003

Binder, A. M., Corvalan, C., Mericq, V., Pereira, A., Santos, J. L., Horvath, S., … Michels, K. B. (2018). Faster ticking rate of the epigenetic clock is associated with faster pubertal development in girls. Epigenetics, 13(1), 85–94. doi: 10.1080/15592294.2017.1414127

Bonen, L. (2008). Cis-and trans-splicing of group II introns in plant mitochondria. Mitochondrion, 8(1), 26–34. doi: 10.1016/j.mito.2007.09.005

Bors, E. K., Baker, C. S., Wade, P. R., O’Neill, K. B., Shelden, K. E. W., Thompson, M. J., … Horvath, S. (2021). An epigenetic clock to estimate the age of living beluga whales. Evolutionary Applications, 14(5), 1263–1273. doi: https://doi.org/10.1111/eva.13195

Chaudhary, S., Jabre, I., & Syed, N. H. (2021). Epigenetic differences in an identical genetic background modulate alternative splicing in A. thaliana. Genomics, 113(6), 3476–3486. doi: 10.1016/j.ygeno.2021.08.006

Christensen, B. C., Houseman, E. A., Marsit, C. J., Zheng, S., Wrensch, M. R., Wiemels, J. L., … Kelsey, K. T. (2009). Aging and Environmental Exposures Alter Tissue-Specific DNA Methylation Dependent upon CpG Island Context. PLOS Genetics, 5(8), e1000602. doi: 10.1371/journal.pgen.1000602

Clason, T. R. (1989). Early Growth Enhancement Increases Loblolly Pine Rotation Yields. Southern Journal of Applied Forestry, 13(2), 94–99. doi: 10.1093/sjaf/13.2.94

Coyle, D. R., Aubrey, D. P., & Coleman, M. D. (2016). Growth responses of narrow or broad site adapted tree species to a range of resource availability treatments after a full harvest rotation. Forest Ecology and Management, 362, 107–119. doi: 10.1016/j.foreco.2015.11.047

Cunningham, K. (n.d.). Managing Loblolly Pine Stands…from A to Z. 6.

Dubin, M. J., Zhang, P., Meng, D., Remigereau, M.-S., Osborne, E. J., Paolo Casale, F., … Nordborg, M. (2015). DNA methylation in Arabidopsis has a genetic basis and shows evidence of local adaptation. ELife, 4, e05255. doi: 10.7554/eLife.05255

Dubrovina, A. S., & Kiselev, K. V. (2016). Age-associated alterations in the somatic mutation and DNA methylation levels in plants. Plant Biology, 18(2), 185–196. doi: 10.1111/plb.12375

Flatt, T., & Partridge, L. (2018). Horizons in the evolution of aging. BMC Biology, 16(1), 1–13.

Fox, T. R., Lee Allen, H., Albaugh, T. J., Rubilar, R., & Carlson, C. A. (2007). Tree nutrition and forest fertilization of pine plantations in the southern United States. Southern Journal of Applied Forestry, 31(1), 5–11.

Friedman, J., Hastie, T., & Tibshirani, R. (2010). Regularization Paths for Generalized Linear Models via Coordinate Descent. Journal of Statistical Software, 33(1), 1–22.

Gehring, M., & Henikoff, S. (2007). DNA methylation dynamics in plant genomes. Biochimica et Biophysica Acta (BBA) - Gene Structure and Expression, 1769(5), 276–286. doi: 10.1016/j.bbaexp.2007.01.009

Gendrel, A.-V., Lippman, Z., Yordan, C., Colot, V., & Martienssen, R. (2002). Dependence of Heterochromatic Histone H3 Methylation Patterns on the Arabidopsis Gene DDM1. Science (New York, N.Y.), 297, 1871–1873. doi: 10.1126/science.1074950

Ghoshal, K., Majumder, S., Datta, J., Motiwala, T., Bai, S., Sharma, S. M., … Jacob, S. T. (2004). Role of Human Ribosomal RNA (rRNA) Promoter Methylation and of Methyl-CpG-binding Protein MBD2 in the Suppression of rRNA Gene Expression *. Journal of Biological Chemistry, 279(8), 6783–6793. doi: 10.1074/jbc.M309393200

Hamlat, E. J., Prather, A. A., Horvath, S., Belsky, J., & Epel, E. S. (2021). Early life adversity, pubertal timing, and epigenetic age acceleration in adulthood. Developmental Psychobiology, 63(5), 890–902. doi: 10.1002/dev.22085

Hannum, G., Guinney, J., Zhao, L., Zhang, L., Hughes, G., Sadda, S., … Zhang, K. (2013). Genome-wide Methylation Profiles Reveal Quantitative Views of Human Aging Rates. Molecular Cell, 49(2), 359–367. doi: 10.1016/j.molcel.2012.10.016

Hastie T., Tibshirani, R., Narasimhan, B., & Chu, G. (2021). impute: impute: Imputation for microarray data. R package version 1.68.0.

Horvath, S. (2013). DNA methylation age of human tissues and cell types. Genome Biology, 14(10), 1–20.

Horvath, S., & Raj, K. (2018). DNA methylation-based biomarkers and the epigenetic clock theory of ageing. Nature Reviews Genetics, 19(6), 371–384.

Hsieh, W.-Y., Liao, J.-C., Chang, C.-Y., Harrison, T., Boucher, C., & Hsieh, M.-H. (2015). The SLOW GROWTH3 Pentatricopeptide Repeat Protein Is Required for the Splicing of Mitochondrial NADH Dehydrogenase Subunit7 Intron 2 in Arabidopsis. Plant Physiology, 168(2), 490–501. doi: 10.1104/pp.15.00354

Hu, Y., Morota, G., Rosa, G. J. M., & Gianola, D. (2015). Prediction of Plant Height in Arabidopsis thaliana Using DNA Methylation Data. Genetics, 201(2), 779–793. doi: 10.1534/genetics.115.177204

Jeddeloh, J. A., Stokes, T. L., & Richards, E. J. (1999). Maintenance of genomic methylation requires a SWI2/SNF2-like protein. Nature Genetics, 22(1), 94–97. doi: 10.1038/8803

Jiang, C., Mithani, A., Belfield, E. J., Mott, R., Hurst, L. D., & Harberd, N. P. (2014). Environmentally responsive genome-wide accumulation of de novo Arabidopsis thaliana mutations and epimutations. Genome Research, 24(11), 1821–1829. doi: 10.1101/gr.177659.114

Jung, M., & Pfeifer, G. P. (2015). Aging and DNA methylation. BMC Biology, 13(1), 7. doi: 10.1186/s12915-015-0118-4

Kabacik, S., Horvath, S., Cohen, H., & Raj, K. (2018). Epigenetic ageing is distinct from senescence-mediated ageing and is not prevented by telomerase expression. Aging (Albany NY), 10(10), 2800–2815. doi: 10.18632/aging.101588

Kellner, K. F., & Swihart, R. K. (2016). Timber harvest and drought interact to impact oak seedling growth and survival in the Central Hardwood Forest. Ecosphere, 7(10), e01473. doi: 10.1002/ecs2.1473

Krueger, F., & Andrews, S. R. (2011). Bismark: A flexible aligner and methylation caller for Bisulfite-Seq applications. Bioinformatics, 27(11), 1571–1572. doi: 10.1093/bioinformatics/btr167

Kuhn, M. (2015). Caret: Classification and regression training. Astrophysics Source Code Library, ascl-1505.

Law, J. A., & Jacobsen, S. E. (2010). Establishing, maintaining and modifying DNA methylation patterns in plants and animals. Nature Reviews Genetics, 11(3), 204–220.

Lawrence, M., Huber, W., Pagès, H., Aboyoun, P., Carlson, M., Gentleman, R., … Carey, V. J. (2013). Software for Computing and Annotating Genomic Ranges. PLOS Computational Biology, 9(8), e1003118. doi: 10.1371/journal.pcbi.1003118

Lemaître, J.-F., Rey, B., Gaillard, J.-M., Régis, C., Gilot, E., Pellerin, M., … Horvath, S. (2020). Epigenetic clock and DNA methylation studies of roe deer in the wild (p. 2020.09.21.306613). doi: 10.1101/2020.09.21.306613

Levine, M. E., Lu, A. T., Quach, A., Chen, B. H., Assimes, T. L., Bandinelli, S., … Horvath, S. (2018). An epigenetic biomarker of aging for lifespan and healthspan. Aging (Albany NY), 10(4), 573–591. doi: 10.18632/aging.101414

Li, B., Mckeand, S., & Weir, R. (1999). Tree improvement and sustainable forestry—Impact of two cycles of loblolly pine breeding in the USA. For. Genet., 6, 229–234.

Li, H., Handsaker, B., Wysoker, A., Fennell, T., Ruan, J., Homer, N., … 1000 Genome Project Data Processing Subgroup. (2009). The Sequence Alignment/Map format and SAMtools. Bioinformatics, 25(16), 2078–2079. doi: 10.1093/bioinformatics/btp352

Longo, G. P., & Scandalios, J. G. (1969). Nuclear gene control of mitochondrial malic dehydrogenase in maize. Proceedings of the National Academy of Sciences, 62(1), 104–111.

Madeira, F., Park, Y. M., Lee, J., Buso, N., Gur, T., Madhusoodanan, N., … Finn, R. D. (2019). The EMBL-EBI search and sequence analysis tools APIs in 2019. Nucleic Acids Research, 47(W1), W636–W641.

Malek, O., & Knoop, V. (1998). Trans-splicing group II introns in plant mitochondria: The complete set of cis-arranged homologs in ferns, fern allies, and a hornwort. RNA, 4(12), 1599–1609. doi: 10.1017/S1355838298981262

Mayne, B., Berry, O., Davies, C., Farley, J., & Jarman, S. (2019). A genomic predictor of lifespan in vertebrates. Scientific Reports, 9(1), 17866. doi: 10.1038/s41598-019-54447-w

Mayne, B., Korbie, D., Kenchington, L., Ezzy, B., Berry, O., & Jarman, S. (2020). A DNA methylation age predictor for zebrafish. Aging (Albany NY), 12(24), 24817–24835. doi: 10.18632/aging.202400

Medlyn, B., Barrett, D., Landsberg, J., Sands, P., & Clement, R. (2003). Corrigendum to: Conversion of canopy intercepted radiation to photosynthate: a review of modelling approaches for regional scales. Functional Plant Biology, 30(7), 829–829. doi: 10.1071/fp02088_co

Meek, M. H., & Larson, W. A. (2019). The future is now: Amplicon sequencing and sequence capture usher in the conservation genomics era. Molecular Ecology Resources, 19(4), 795–803. doi: 10.1111/1755-0998.12998

Meer, M. V., Podolskiy, D. I., Tyshkovskiy, A., & Gladyshev, V. N. (2018). A whole lifespan mouse multi-tissue DNA methylation clock. Elife, 7, e40675.

Neale, D. B., Wegrzyn, J. L., Stevens, K. A., Zimin, A. V., Puiu, D., Crepeau, M. W., … Langley, C. H. (2014). Decoding the massive genome of loblolly pine using haploid DNA and novel assembly strategies. Genome Biology, 15(3), R59. doi: 10.1186/gb-2014-15-3-r59

Ng, H.-H., & Adrian, B. (1999). DNA methylation and chromatin modification. Current Opinion in Genetics & Development, 9(2), 158–163. doi: 10.1016/S0959-437X(99)80024-0

Ong-Abdullah, M., Ordway, J. M., Jiang, N., Ooi, S.-E., Kok, S.-Y., Sarpan, N., … Martienssen, R. A. (2015). Loss of Karma transposon methylation underlies the mantled somaclonal variant of oil palm. Nature, 525(7570), 533–537. doi: 10.1038/nature15365

Parrott, B. B., & Bertucci, E. M. (2019). Epigenetic aging clocks in ecology and evolution. Trends in Ecology & Evolution, 34(9), 767–770.

Partridge, L., & Gems, D. (2002). Mechanisms of aging: Public or private? Nature Reviews Genetics, 3(3), 165–175.

Perls, T., Kunkel, L., & Puca, A. (2002). The genetics of aging. Current Opinion in Genetics & Development, 12(3), 362–369.

Perna, L., Zhang, Y., Mons, U., Holleczek, B., Saum, K.-U., & Brenner, H. (2016). Epigenetic age acceleration predicts cancer, cardiovascular, and all-cause mortality in a German case cohort. Clinical Epigenetics, 8(1), 64. doi: 10.1186/s13148-016-0228-z

Probst, A. V., & Mittelsten Scheid, O. (2015). Stress-induced structural changes in plant chromatin. Current Opinion in Plant Biology, 27, 8–16. doi: 10.1016/j.pbi.2015.05.011

Quinlan, A. R., & Hall, I. M. (2010). BEDTools: A flexible suite of utilities for comparing genomic features. Bioinformatics, 26(6), 841–842. doi: 10.1093/bioinformatics/btq033

R Core Team (2021). R: A language and environment for statistical computing. R Foundation for Statistical Computing, Vienna, Austria. https://www.R-project.org/.

Raddatz, G., Arsenault, R. J., Aylward, B., Whelan, R., Böhl, F., & Lyko, F. (2021). A chicken DNA methylation clock for the prediction of broiler health. Communications Biology, 4(1), 1–8. doi: 10.1038/s42003-020-01608-7

Revelle, W. (2019). psych: Procedures for psychological, psychometric, and personality research. Northwestern University, Evanston, Illinois. R package version 1.9. 12. URL http://CRAN.R-Project.Org/Package=Psych.

Richardson, B. (2003). Impact of aging on DNA methylation. Ageing Research Reviews, 2(3), 245–261. doi: 10.1016/S1568-1637(03)00010-2

Russell, J., & Zomerdijk, J. C. B. M. (2005). RNA-polymerase-I-directed rDNA transcription, life and works. Trends in Biochemical Sciences, 30(2), 87–96. doi: 10.1016/j.tibs.2004.12.008

Ryan, C. P. (2021). “Epigenetic clocks”: Theory and applications in human biology. American Journal of Human Biology, 33(3), e23488. doi: 10.1002/ajhb.23488

Ryan, C. P., Hayes, M. G., Lee, N. R., McDade, T. W., Jones, M. J., Kobor, M. S., … Eisenberg, D. T. A. (2018). Reproduction predicts shorter telomeres and epigenetic age acceleration among young adult women. Scientific Reports, 8(1), 11100. doi: 10.1038/s41598-018-29486-4

Samuelson, L., Stokes, T., Cooksey, T., & McLemore III, P. (2001). Production efficiency of loblolly pine and sweetgum in response to four years of intensive management. Tree Physiology, 21(6), 369–376.

Schulze, M. D. (2003). Ecology and behavior of nine timber tree species in Pará, Brazil: Links between species life history and forest management and conservation. The Pennsylvania State University.

Shahryary, Y., Symeonidi, A., Hazarika, R. R., Denkena, J., Mubeen, T., Hofmeister, B., … Johannes, F. (2020). AlphaBeta: Computational inference of epimutation rates and spectra from high-throughput DNA methylation data in plants. Genome Biology, 21(1), 260. doi: 10.1186/s13059-020-02161-6

Sharma, R., & Patnaik, S. K. (1982). Differential Regulation of Malate Dehydrogenase Isoenzymes by Hydrocortisone in the Liver and Brain of Aging Rats. Development, Growth & Differentiation, 24(5), 501–505. doi: 10.1111/j.1440-169X.1982.00501.x

Simpkin, A. J., Suderman, M., Gaunt, T. R., Lyttleton, O., McArdle, W. L., Ring, S. M., … Relton, C. L. (2015). Longitudinal analysis of DNA methylation associated with birth weight and gestational age. Human Molecular Genetics, 24(13), 3752–3763. doi: 10.1093/hmg/ddv119

Slotkin, R. K., & Martienssen, R. (2007). Transposable elements and the epigenetic regulation of the genome. Nature Reviews Genetics, 8(4), 272–285.

Stiff, C. T., & Stansfield, W. F. (2003). Thinning guidelines for loblolly pine plantations in eastern Texas based on alternative management criteria. 12th Biennial Southern Silvicultural Research Conference, 323.

Stubbs, T. M., Bonder, M. J., Stark, A.-K., Krueger, F., Bolland, D., Butcher, G., … BI Ageing Clock Team. (2017). Multi-tissue DNA methylation age predictor in mouse. Genome Biology, 18(1), 68. doi: 10.1186/s13059-017-1203-5

Suzuki, M. M., & Bird, A. (2008). DNA methylation landscapes: Provocative insights from epigenomics. Nature Reviews Genetics, 9(6), 465–476. doi: 10.1038/nrg2341

Sweetman, C., Waterman, C. D., Rainbird, B. M., Smith, P. M. C., Jenkins, C. D., Day, D. A., & Soole, K. L. (2019). AtNDB2 Is the Main External NADH Dehydrogenase in Mitochondria and Is Important for Tolerance to Environmental Stress1 [OPEN]. Plant Physiology, 181(2), 774–788. doi: 10.1104/pp.19.00877

Takuno, S., Ran, J.-H., & Gaut, B. S. (2016). Evolutionary patterns of genic DNA methylation vary across land plants. Nature Plants, 2(2), 1–7. doi: 10.1038/nplants.2015.222

Tomaz, T., Bagard, M., Pracharoenwattana, I., Lindén, P., Lee, C. P., Carroll, A. J., … Millar, A. H. (2010). Mitochondrial Malate Dehydrogenase Lowers Leaf Respiration and Alters Photorespiration and Plant Growth in Arabidopsis[W][OA]. Plant Physiology, 154(3), 1143–1157. doi: 10.1104/pp.110.161612

van der Graaf, A., Wardenaar, R., Neumann, D. A., Taudt, A., Shaw, R. G., Jansen, R. C., … Johannes, F. (2015). Rate, spectrum, and evolutionary dynamics of spontaneous epimutations. Proceedings of the National Academy of Sciences, 112(21), 6676–6681. doi: 10.1073/pnas.1424254112

Weidner, C. I., Lin, Q., Koch, C. M., Eisele, L., Beier, F., Ziegler, P., … Wagner, W. (2014). Aging of blood can be tracked by DNA methylation changes at just three CpG sites. Genome Biology, 15(2), R24. doi: 10.1186/gb-2014-15-2-r24

Weiss, H., Friedrich, T., Hofhaus, G., & Preis, D. (1992). The respiratory-chain NADH dehydrogenase (complex I) of mitochondria. In P. Christen & E. Hofmann (Eds.), EJB Reviews 1991 (pp. 55–68). Berlin, Heidelberg: Springer. doi: 10.1007/978-3-642-77200-9_5

Xiao, F.-H., Wang, H.-T., & Kong, Q.-P. (2019). Dynamic DNA Methylation During Aging: A “Prophet” of Age-Related Outcomes. Frontiers in Genetics, 0. doi: 10.3389/fgene.2019.00107

Yao, N., Schmitz, R. J., & Johannes, F. (2021). Epimutations define a fast-ticking molecular clock in plants. Trends in Genetics.

Yudina, R. S. (2012). Malate dehydrogenase in plants: Its genetics, structure, localization and use as a marker. 2012. doi: 10.4236/abb.2012.34053

Zatsepina, O. V., Voit, R., Grummt, I., Spring, H., Semenov, M. V., & Trendelenburg, M. F. (1993). The RNA polymerase I-specific transcription initiation factor UBF is associated with transcriptionally active and inactive ribosomal genes. Chromosoma, 102(9), 599–611. doi: 10.1007/BF00352307

Zhang, H., Lang, Z., & Zhu, J.-K. (2018). Dynamics and function of DNA methylation in plants. Nature Reviews Molecular Cell Biology, 19(8), 489–506. doi: 10.1038/s41580-018-0016-z

Zheng, Y., Joyce, B. T., Colicino, E., Liu, L., Zhang, W., Dai, Q., … Hou, L. (2016). Blood Epigenetic Age may Predict Cancer Incidence and Mortality. EBioMedicine, 5, 68–73. doi: 10.1016/j.ebiom.2016.02.008

Zilberman, D. (2008). The evolving functions of DNA methylation. Current Opinion in Plant Biology, 11(5), 554–559. doi: 10.1016/j.pbi.2008.07.004

